# Characterising the performance of an antibiotic resistance prediction tool, gnomonicus, using a diverse testset of 2,663 *Mycobacterium tuberculosis* samples

**DOI:** 10.1101/2024.11.08.622466

**Authors:** Jeremy Westhead, Catriona S Baker, Marc Brouard, Matthew Colpus, Bede Constantinides, Alexandra Hall, Jeff Knaggs, Marcela Lopes Alves, Ruan Spies, Hieu Thai, Sarah Surrall, Kumeren Govender, Timothy EA Peto, Derrick W Crook, Shaheed V Omar, Robert Turner, Philip W Fowler

## Abstract

Tuberculosis remains a global health problem. Making it easier and quicker to identify which antibiotics an infection is likely to be susceptible to will be a key part of the solution. Whilst whole-genome sequencing offers many advantages, the processing of the genetic reads to produce the relevant public health and clinical information is, surprisingly, often the responsibility of the end user which inhibits uptake. Here we characterise how well a freely-available tool we have developed, gnomonicus, predicts the antibiotic resistance profile of a sample (given its variant call file) using our implementation of the second edition of the WHO catalogue of resistance-associated variants (WHOv2). To facilitate this, we have constructed a Diverse Testset of 2,663 publicly-available *M. tuberculosis* samples which have both genetic and drug susceptibility testing (DST) data. We have chosen to apply the catalogue such that our tool will return a result of (i) Fail if there are insufficient reads at a genetic locus associated with resistance, (ii) Unknown if a genetic variant in a resistance gene not listed in the catalogue is encountered and (iii) Resistant if three or more short-reads support the presence of a resistance-associated variant. The last step increases the sensitivity for all 15 antibiotics but only reaches significance in a few in our testset. Comparing our results to those of `TB-Profiler`, an existing tool, highlights the different design choices and demonstrates the performance of both tools on our Diverse Testset is comparable. By only considering high confidence DST results we show that gnomonicus, in combination with our translation of WHOv2, achieves sensitivities and specificities in excess of 95% for both isoniazid and rifampicin.

**Impact Statement:** Whole genome sequencing clinical samples taken from patients with tuberculosis is a potentially fast and accurate method for determining to which antibiotics the infection will be susceptible. Two barriers need to be overcome; the first, which is knowing which mutations are associated with resistance (or not) to a range of antibiotics is well on the way to be solved thanks to the efforts of the World Health Organization (WHO) who have published extensive catalogues containing lists of such mutations. The second barrier is that the processing of the raw genetic files remains a largely manual process overseen by bioinformaticians. Here we describe `gnomonicus`, our open-source AMR prediction tool, and report the performance of our translation of the second edition of the WHO catalogue using a carefully designed publicly-available dataset of 2,663 *M. tuberculosis* samples. We hope that not only will this tool be useful but also that this dataset will be used by other researchers to facilitate comparisons between pipelines, approaches and tools.

**Data Summary:** The attendant GitHub repository^1^ allows `gnomonicus` to be rerun on all 2,663 samples in the Diverse Testset; it therefore includes instructions, the necessary input files (all the variant call files, the version of the WHOv2 catalogue and a link to the H37Rv GenBank file used in this study) and also the output JSON files. The ENA accession numbers for all 2,663 samples (including a bash script to download them) and their corresponding phenotypic drug susceptibility testing results are included with the intention that people can either reproduce our results, or use the same dataset for other analyses. The JSON files output by `TB-Profiler` have also been added for comparison. The repository contains a series of Juypter notebooks containing Python3 code that allows the user to discover and parse the output JSON files from either tool and save the results as data tables. Other notebooks allow the user to reproduce all the analysis underlying this work, including reproducing the figures and many of the tables.

## Introduction

In 2023 just under 11 million people became ill with tuberculosis (TB) worldwide and 1.25 million died^2^. The aetiological agent, *M. tuberculosis*, requires treating with multiple antibiotics. Drug-susceptible *M. tuberculosis* is routinely treated with four antibiotics: rifampicin, isoniazid, pyrazinamide and ethambutol. Like other bacterial pathogens, resistance to all antibiotics has now been observed and multi-drug resistant (MDR) TB is defined as infections resistant to both rifampicin and isoniazid – these require alternative treatment, such as the BPaLM (bedaquline, protonamid, linezolid and moxifloxacin) regimen that was recommended by the WHO in 2022^3^. MDR-TB is increasingly recognised as a global health concern and consequently rifampicin-resistant *M. tuberculosis* was added to the WHO Bacterial Priority Pathogens List in 2024^4^. Positive identification of *M. tuberculosis* complex in a clinical sample and subsequent phenotypic drug susceptibility testing (pDST) are important steps in treating this disease.

Due to the slow growth rate of *M. tuberculosis* culture-based pDST takes weeks; whole genome sequencing (WGS) is an attractive alternative since it is faster, potentially more accurate and also yields epidemiological information. Over the last eight years^5^, many high income countries have adopted WGS for *M. tuberculosis*. Our ability to accurately predict the antibiogram of a sample from its genetics has been boosted in recent years by the systematic sequencing and drug susceptibility testing of large numbers of clinical samples by projects such as the Comprehensive Resistance Prediction for Tuberculosis: an International Consortium (CRyPTIC). CRyPTIC collected over 15,000 clinical samples via 14 laboratories based in 11 countries^6^. This and other publicly available datasets enabled the World Health Organization to release in 2021 the first catalogue of mutations in *M. tuberculosis* complex associated with drug resistance^7^. A second edition was released in late 2023^8^, which we will call WHOv2. Both editions take the form of a traditional text-based report (which contains some expert rules) and an accompanying detailed Excel spreadsheet (with more rules), with the second edition also including a variant call file. Each edition used a single dataset to both infer the association of genetic variants with antibiotic resistance and estimate the performance of the resulting catalogue. It is challenging to apply either catalogue to the WGS data of a clinical sample since no computational tool was released with either catalogue nor was an annotated test dataset released that would have simplified tool development.

In this paper we shall describe a tool, `gnomonicus`, we have developed that ingests a resistance catalogue, a sample variant call file and the GenBank file of the reference genome and outputs the mutations detected and their predicted effects on a range of antibiotics. Since no validation dataset is publicly available, we have also built a Diverse Testset of 2,663 *M. tuberculosis* samples that undergone both whole gene sequencing and pDST. This will let us calculate the performance of our implementation of the WHOv2 catalogue and compare it to the stated performance. Finally, we compare our results with those of TB-Profiler^9^ which is the best known third-party tool that performs a similar function.

## Materials and Methods

### Sample selection

A total of 11,887 samples were identified from v2.1.2 of the publicly available CRyPTIC dataset^10^ that (i) had been whole genome sequenced using short-read (Illumina) technologies and (ii) had minimum inhibitory concentrations (MICs) to 13 different antibiotics measured using a bespoke 96-well broth microdilution (BMD) plate. Each sample was sequenced and incubated on a BMD plate as described previously^6,11^. In addition to the MICs measured visually by the laboratory scientist, photographs of each plate were image processed^12^ and classified by at least 11 volunteers as part of a citizen science project^13^. All MICs where two or three of these independent measurements agreed were annotated as high confidence MICs – these are assumed to have reduced measurement error. Two plate designs were used (UKMYC5 & UKMYC6), each of which included 13 antibiotics: amikacin, bedaquline, clofazimine delamanid, ethambutol, ethionamide, isoniazid, kanamycin, levofloxacin, linezolid, moxifloxacin, rifabutin and rifampicin. All MICs were converted to a binary `Resistant`/`Susceptible` result using a set of research ECOFFs^11^. All 13 antibiotics bar rifabutin are included in the WHOv2 catalogue. The WHOv2 catalogue also includes three drugs not present on the plates: pyrazinamide, capromycin and streptomycin. We therefore also identified a further 10,606 publicly available samples from the CRyPTIC dataset^10^ which had at least one binary phenotype of these drugs measured via the MGIT960 system. In practice, many of these samples will have been tested using both pDST methods, however, to avoid any potential inconsistencies introduced by combining the results from different methods, each sample is associated with only one pDST method.

### Construction of the Diverse Testset

The Diverse Testset has two competing aims: (i) have as close to 50% resistance / 50% susceptibility for all drugs to maximise resolution whilst also (ii) being as small as possible to enable rapid, repeated testing. The difficulty being, of course, that each sample selected has phenotypic information for more than one drug which makes it difficult to achieve the first criterion. Finally there is a risk of introducing bias if all samples have some degree of resistance. We therefore arbitrarily decided to create an initial dataset of 1,000 samples with phenotypes for the 13 drugs on the UKMYC plate designs ensuring that 200 of these were pan-susceptible.

Since the CRyPTIC project collected very few samples resistant to the new and repurposed drugs, we first selected all 284 samples which were assessed as resistant to one or more of bedaquline (n=65), linezolid (117) and delamanid (140). Additional samples were then randomly selected if they (i) were resistant to the remaining drug with the fewest resistant samples, (ii) were not resistant to any drugs that had already reached 50% penetration in the dataset and (iii) were drawn from the remaining 100 samples with the greatest number of high confidence MICs. This process was repeated until 800 samples had been chosen whereupon a further 200 pan-susceptible samples were chosen from the 1,000 pan-susceptible samples with the greatest number of high confidence MICs.

Despite our efforts, there are fewer than 250 resistant samples for bedaquline, linezolid, and delamanid in the UKMYC dataset (Table 1). We therefore repeated the process of iteratively selecting individual samples on the dataset of 10,606 samples with MGIT pDST data, the main differences being (i) we only considered pyrazinamide, capreomycin, streptomycin, linezolid and bedaquline and (ii) each sample had pDST results for a variable number of these drugs and therefore we tried to maximise the number of pDST results per sample. After de-duplication with the UKMYC dataset and verification that all samples had raw genetic (FASTQ) files available in the European Nucleotide Archive (ENA), this led to a dataset of 1,663 samples. The number of pDST results per sample (Table S1) varied between one and four with two being the most common. The aggregated dataset of 2,663 samples therefore contains at least 250 resistant samples for all drugs (Table 1), except linezolid (n=117) and delamanid (140). It is important to note that the process to produce the Diverse Testset was designed to satisfy the aims stated above but does not ensure a diversity in resistance mechanisms.

**Table 1:** The proportion of resistance by drug in the 1,000 UKMYC and 1,663 MGIT samples. Due to having the highest prevalence of resistance in the CRyPTIC dataset, only isoniazid reached 50% resistance. A key difference between the two datasets is that each UKMYC samples has an MIC and thence a binary phenotype for 13 drugs whereas the MGIT samples have between one and four binary phenotypes, with two being the most common (702 samples, Table S1). As each UKMYC plate was read by three independent methods, all MICs where at least two methods agree are assumed to be high confidence.

| Antibiotic | pDST method | %Resistant | R | S | TOTAL | %High confidence |
| --- | --- | --- | --- | --- | --- | --- |
| Bedaquiline | UKMYC | 6.5 | 65 | 935 | 1000 | 85.1 |
| Linezolid | UKMYC | 11.7 | 117 | 883 | 1000 | 71.9 |
| Delamanid | UKMYC | 14.0 | 140 | 860 | 1000 | 74.9 |
| Ethambutol | UKMYC | 28.8 | 288 | 712 | 1000 | 76.7 |
| Moxifloxacin | UKMYC | 28.8 | 288 | 712 | 1000 | 73.2 |
| Clofazimine | UKMYC | 28.8 | 288 | 712 | 1000 | 67.6 |
| Amikacin | UKMYC | 28.8 | 288 | 712 | 1000 | 82.5 |
| Ethionamide | UKMYC | 28.8 | 288 | 712 | 1000 | 86.3 |
| Kanamycin | UKMYC | 28.9 | 289 | 711 | 1000 | 88.0 |
| Levofloxacin | UKMYC | 32.3 | 323 | 677 | 1000 | 79.0 |
| Rifampicin | UKMYC | 47.6 | 476 | 524 | 1000 | 88.0 |
| Isoniazid | UKMYC | 50.0 | 500 | 500 | 1000 | 90.6 |
| Linezolid | MGIT | 13.5 | 105 | 675 | 780 | - |
| Streptomycin | MGIT | 41.0 | 252 | 362 | 614 | - |
| Capreomycin | MGIT | 30.8 | 256 | 575 | 831 | - |
| Pyrazinamide | MGIT | 46.4 | 291 | 336 | 627 | - |
| Bedaquiline | MGIT | 50.9 | 394 | 380 | 774 | - |

### Identifying the variants in the Diverse Testset using EIT GPAS

For simplicity, all samples were processed using the Mycobacterial pipeline deployed in the EIT Global Pathogen Analysis Service (GPAS, https://gpas.global). The pipeline used is described in more detail in the Supplement. Samples were uploaded using its command line interface (CLI). This ensured reads matching the human genome were removed using the `hostile`^14^ algorithm prior to upload. Upon arrival in the cloud all samples undergo a second round of decontamination with `hostile`^14^ then poor quality/short reads are discarded before the number of reads belonging to the Mycobacterium genus is assessed using `kraken2` (v2.1.3) in conjunction with the `Standard` index from June 2023^15^. If a sample has over 10,000 Mycobacterial (and unclassified) short reads it is progressed. These reads are then competitively mapped to a curated list of 186 Mycobacterial genomes using `minimap2` (v2.24-r1122)^16^ to provide fine-grained speciation. Lastly, in the case of complexes the reads are then examined by `mykrobe` (v0.13.0)^17^ which classifies reads down to the level of lineage (in the case of *Mycobacterium tuberculosis* complex) or subspecies (for *Mycobacterium avium* complex and *Mycobacterium abscessus* complex). Reads are then mapped to version 3 of the *M. tuberculosis* H37Rv reference genome^18–20^ (NC 000962.3) using `clockwork` (v0.12.3)^21^ which in turn uses `minimap2`^16^ to build a pile-up and then both `samtools` (v1.15.1)^22^ and `cortex`^23^ to call variants; the former being better at identifying SNPs and the latter insertions and deletions and `minos`^24^ adjudicates when there is overlap. `Clockwork` was used by the CRyPTIC project^6^ and also for all variant calling in the first edition of the WHO catalogue of resistance-associated variants^7^. Clockwork calls a genetic variant if it is supported by 90% or more of the reads. The variant call files for all 2,663 samples were downloaded and stored in the attendant GitHub repository^1^ for later processing by `gnomonicus`.

### Translation of the second edition of the WHO catalogue of resistance-associated variants in *M. tuberculosis* and resistance prediction

As mentioned, WHOv2 comprises three artefacts: no single artefact contains all the rules in the catalogue and only the Excel and VCF files are parsable by computer code. The Excel file adopts the HGVS nomen-clature for describing genetic variants but this is unable to encode some of the broader rules found in the catalogue such as *“any frameshift in gene X”* which is a key component of the new Loss of Function rules. The CRyPTIC project^11^ developed a grammar for antimicrobial resistance catalogues (GARC) specifically for this purpose and we therefore translated the Excel file into the GARC format^25^ that can be read and understood by `gnomonicus`^26^. A formal definition of the GARC grammar is provided via its Backus-Naur Form in the Supplement. Code changes were required to incorporate the new epistatic rules introduced in WHOv2. Our catalogue maps Groups 1 & 2 onto `Resistant` (`R`) and Groups 4 & 5 onto `Susceptible` (`S`) as suggested by the WHO, with Group 3 being labelled `Unknown` (`U`).

There are three important differences in our implementation of WHOv2; our catalogue will report any mutation in a gene known to be associated with resistance but not in the catalogue as `Unknown`. If there are two or fewer reads at a genetic locus known to be associated with resistance our catalogue will return a result of `Fail` since there is not enough information to know if the sample is resistant or not, and it probably needs re-sequencing. Neither scenario is described by WHOv2 and therefore both would be predicted susceptible without these rules. Finally, we call any resistance-associated variant listed in the catalogue if it is supported by three or more reads, regardless of how many other (usually wild-type) reads are also found at that locus. Minor alleles with resistance-associated variants are therefore detected; by contrast 75% of reads were required to support a genetic variant for it to be considered for the WHOv2 catalogue and there is no guidance on what threshold should be applied when detecting variants listed in WHOv2.

### Predicting the antibiogram of a sample using gnomonicus

Our translation of the WHOv2 catalogue^27^, along with the variant call file produced by `clockwork` and version 3 of the reference H37Rv genome (as a GenBank file) are then ingested by `gnomonicus` (v3.0.10)^26^ which produces lists of which genetic variants are detected in the sample (translated into amino acid mutations where appropriate), their predicted effects on antibiotics as specified by the catalogue and finally the resulting predicted antibiogram. The resulting javascript object notation (JSON) files, along with the command used to generate them, are stored in the attendant repository^1^. Since the pipeline deployed in GPAS also uses `gnomonicus`, we also downloaded an output file for each sample using the same CLI that handled uploading; this contains general information about the sample. All 5,326 output files are stored in the attendant GitHub repository^1^ as described in the Data Summary.

### Comparison with TB-Profiler

All 2,663 samples were also processed by `TB-Profiler` (v6.6.5)^9^, an established bioinformatic tool. It uses `GATK`^28^ for variant calling, rather than Clockwork. `TB-Profiler` produces a JSON file containing a list of detected mutations and their predicted effects on specific antibiotics. It not only implements its own version of WHOv2 but also includes additional rules for identifying resistance derived from the scientific literature. Unlike our GARC grammar, it has no `Fail` rule and only classifies mutations if they are listed in its catalogue. Hence mutations in resistance genes that are not in the catalogue will be predicted to be `Susceptible`. Five of the 2,663 samples failed to return a result and these were excluded from the subsequent analysis. The resulting output JSON files for the remaining 2,658 samples are included in the attendant GitHub repository^1^, as is code to aggregate all the results into data tables, thereby enabling direct comparison with the results of `gnomonicus`.

## Results

### Some antibiotics performed slightly better, some slightly worse when we applied a Simple implementation of the WHOv2 catalogue

Since the WHOv2 catalogue does not provide any guidance on whether to detect minor alleles or what to do if there are insufficient reads at a loci associated with resistance, we began by ignoring both of these effects and simply calculated the performance on our Diverse Testset – we call this the Simple implementation (Fig. 1, S1 and Table 2, S2). We found five antibiotics had sensitivities within ±1.5% (rifampicin, isoniazid, streptomycin, amikacin, kanamycin, Fig. 1) of that reported by the WHO, two antibiotics (pyrazinamide, ethambutol) had higher sensitivities (*>*1.5%) and the remaining eight drugs (capreomycin, bedaquline, linezolid, moxifloxacin, levofloxacin, clofazimine, delamanid, ethionamide) all recorded lower sensitivities (*<* 1.5%) than seen on the WHOv2 Training Dataset. The variation in specificity was lower: two antibiotics (ethionamide, kanamycin) recorded higher specificities (*>* 1.5%) than reported for WHOv2 with one drug (ethambutol) lower. Comparing our results to those published by the WHOv2 on its own Training Dataset^8^ is unfair since the datasets are different but at least gives us some idea of the performance, although we cannot conclude what is responsible for the observed differences.

**Table 2:** The performance of the WHOv2 catalogue on the Diverse Testset of 2,663 *M. tuberculosis* samples. Fails+Minors includes rules that will Fail a drug if there are two or fewer reads at a locus associated with resistance but will call Resistant if there are three or more reads at the same locus. Simple excludes these rules. Fails+Minors (High confidence) applies the same catalogue, but to a subset of the Diverse Testset with only MICs supported by two or more independent reading methods concur (high confidence). This is only possible for antibiotics with UKMYC pDST data – antibiotics marked with an asterisk used simple binary MGIT pDST and therefore have no value for this last category. Confidence limits were estimated by bootstrapping.

|  | Simple |  | Fails+Minors |  | Fails+Minor<br>(High confidence) |  |
| --- | --- | --- | --- | --- | --- | --- |
|  | sensitivity | specificity | sensitivity | specificity | sensitivity | specificity |
| Isoniazid | 91.4 ±0.5 | 95.3 ±0.4 | 92.8 ±0.5 | 94.4 ±0.4 | 95.6 ±0.4 | 95.1 ±0.4 |
| Rifampicin | 94.3 ±0.5 | 96.0 ±0.3 | 95.7 ±0.4 | 95.0 ±0.4 | 96.4 ±0.3 | 95.3 ±0.4 |
| Pyrazinamide* | 81.7 ±0.7 | 97.4 ±0.3 | 85.8 ±0.7 | 97.0 ±0.4 |  |  |
| Ethambutol | 85.5 ±0.8 | 84.7 ±0.6 | 87.3 ±0.8 | 83.7 ±0.6 | 90.5 ±0.6 | 81.5 ±0.6 |
| Bedaquiline* | 40.7 ±1.0 | 98.6 ±0.2 | 66.5 ±0.9 | 97.8 ±0.3 |  |  |
| Linezolid | 22.4 ±1.9 | 99.9 ±0.1 | 29.2 ±2.1 | 99.8 ±0.1 | 45.0 ±2.2 | 99.8 ±0.1 |
| Moxifloxacin | 81.4 ±0.8 | 94.6 ±0.3 | 86.7 ±0.7 | 93.9 ±0.3 | 90.5 ±0.7 | 93.2 ±0.4 |
| Levofloxacin | 78.1 ±1.0 | 96.8 ±0.3 | 83.4 ±0.9 | 96.3 ±0.3 | 87.5 ±0.7 | 96.6 ±0.3 |
| Clofazimine | 7.4 ±0.7 | 97.8 ±0.2 | 14.8 ±0.9 | 96.4 ±0.3 | 12.7 ±1.5 | 96.5 ±0.2 |
| Delamanid | 12.1 ±1.2 | 99.9 ±0.0 | 12.9 ±1.2 | 99.9 ±0.0 | 21.4 ±2.0 | 100.0 ±0.0 |
| Amikacin | 71.8 ±1.4 | 99.6 ±0.1 | 75.2 ±1.3 | 99.5 ±0.1 | 83.9 ±0.9 | 99.3 ±0.1 |
| Streptomycin* | 80.8 ±0.7 | 95.6 ±0.3 | 84.2 ±0.8 | 94.6 ±0.4 |  |  |
| Ethionamide | 71.5 ±1.0 | 87.3 ±0.5 | 73.3 ±1.0 | 86.0 ±0.5 | 77.9 ±0.9 | 85.3 ±0.5 |
| Kanamycin | 73.9 ±1.3 | 99.0 ±0.1 | 77.5 ±1.2 | 98.8 ±0.1 | 85.4 ±0.8 | 98.4 ±0.1 |
| Capreomycin* | 64.1 ±1.0 | 98.3 ±0.2 | 76.4 ±0.8 | 98.2 ±0.2 |  |  |

**Figure 1.**
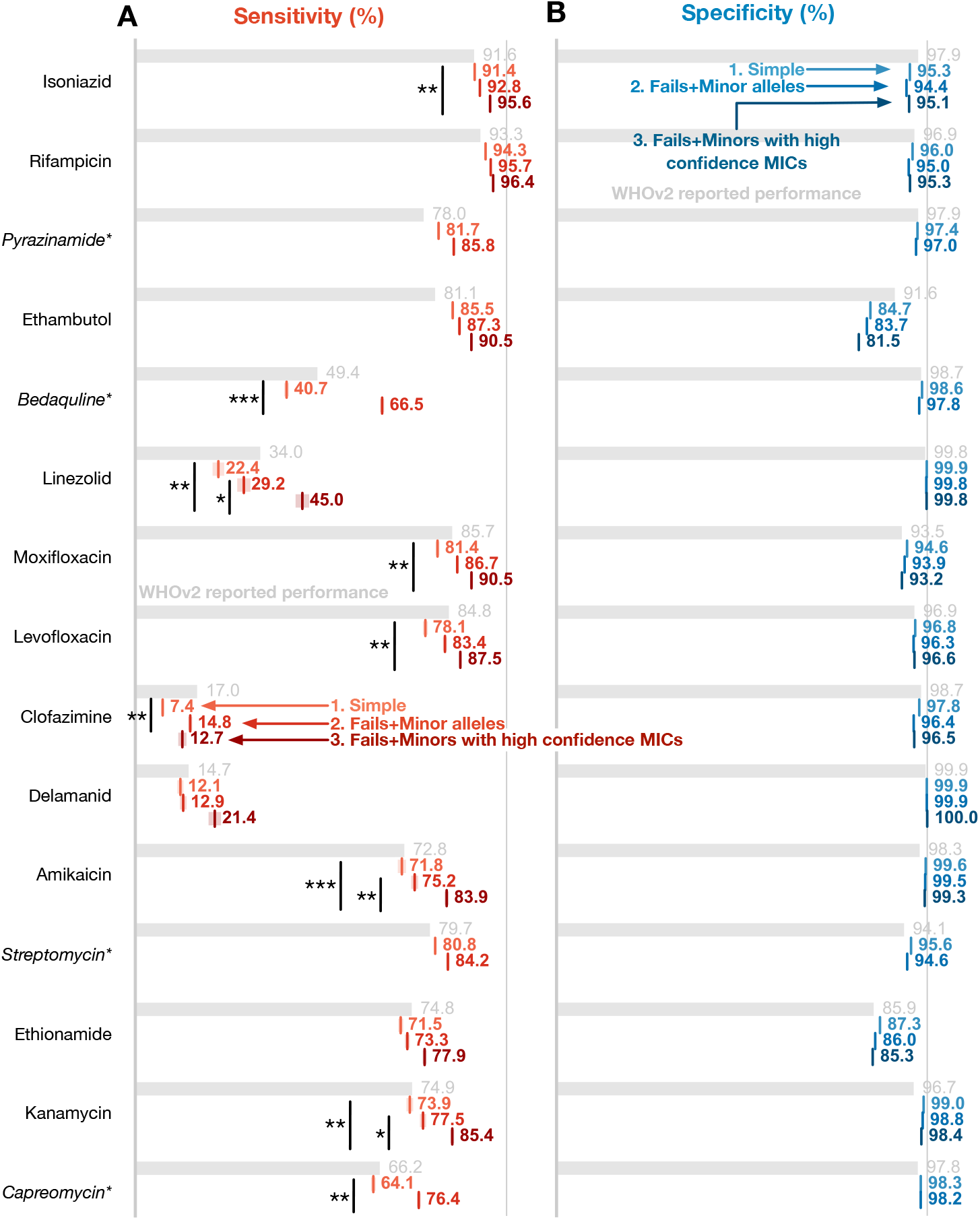
When tested on the 2,663 samples of the Diverse Testset gnomonocius produces comparable AMR prediction performance using our translated version of the WHOv2 catalogue as reported on a much larger dataset^27^. The (**A**) sensitivity and (**B**) susceptibility values for the 15 antibiotics included in the WHOv2 catalogue. The grey bars are the performance reported on the WHO Training Dataset and since are calculated on a different, larger dataset are not directly comparable. Four drugs (marked with an asterisk) have phenotypes from MGIT testing. Our translation of the WHOv2 catalogue is applied in three different ways: (i) a straightforward application, (ii) then allowing resistant minor alleles with three or more reads to be called and, lastly, (ii) only comparing to high confidence MICs in addition to calling minor alleles. Since the latter can only be calculated for the antibiotics with UKMYC MICs, no values can be reported for the four drugs measured by MGIT. Mantel-Haentzsel statistics were calculated for all comparisons. Comparisons with p-values less than 0.05 and 0.01 are annotated with one or two asterisks, respectively.

### Failing samples with insufficient reads at loci associated with resistance is worthwhile but does not improve performance

If there are insufficient reads at a genetic locus one does not know if the nucleotide is the same as the reference, is different resulting in a SNP or even has been deleted, perhaps leading to a frame-shift. If the genetic locus is also one associated with resistance then the likelihood is such that, even in the absence of genetic information, the sample has a reasonable probability of being resistant to the relevant antibiotic. This is a qualitatively different outcome to `R`, `S` or `U` since re-sequencing is likely to result in more reads at that position, allowing a definite result to be returned. In our implementation of the WHOv2 catalogue, any genetic locus in a sample which has two or fewer (including zero) reads but could be a resistance-associated variant (RAV) was described as a `Fail` (`F`).

Adding these rules yielded the Fails implementation. Considering the genetic-based predictions for all 15 drugs using all 2,663 samples yields 194 `Fails` from 39 genetic loci which affected the antibiogram of 66 (2.5%) samples. The majority of `Fail` calls occured in the *rrs* gene (106) followed by *rpoB* (20), *katG* (17), *pncA* (8) and *embB* (9). The resulting changes to the performance of streptomycin, rifampicin, isoniazid, pyrazinamide and ethambutol were small and lay within error and are therefore not shown. Note that the majority of loci in *rrs* with `Fails` did not contribute to amikacin and kanamycin resistance (according to WHOv2) and therefore these antibiotics are, perhaps surprisingly, little affected.

### Identifying minor alleles containing resistance-associated variants improved performance

Thus far all genetic variants have been identified using the default filter in the `clockwork` pipeline which requires 90% of reads to support a variant for it to be called. Note that this threshold is the same as used in WHOv1^7^ but higher than the 75% used in WHOv2^8^. Next, we added rules to our implementation of the WHOv2 catalogue such that any RAV supported by three or more short reads, regardless of how many reads are in support of other (usually wild-type) alleles were called `Resistant`: this is the Fails+Minors implementation. The sensitivity of every antibiotic increased (Fig. 1 and Table 2) with values ranging from +0.8% (streptomycin) to +9.9% (capreomycin). Nine antibiotics experienced an increase of sensitivity of 2% or more (pyrazinamide +4.1%, bedaquline +25.8%, linezolid +6.8%, moxifloxacin +5.3%, levofloxacin +5.3%, clofazimine +7.4%, amikacin +3.4%, kanamycin +3.6% and capreomycin +12.3%) although only the increases seen for bedaquline, clofazimine and capreomycin were statistically significant. Only clofazimine and ethionamide experienced reductions in specificity of more than 1% (1.4% & 1.3%, respectively); neither was statistically significant.

### Only comparing to high confidence pDST results improves the measured performance

There are many reasons why we cannot perfectly predict the antibiogram for each sample; these include sample mislabelling, not all genes or genetic variants having been classified, pDST measurement errors and genetics not being a perfect predictor of phenotype. We are fortunate in that the UKMYC dataset of 1,000 samples has images of all the 96-well plates from when they were read by the laboratory scientist after two weeks incubation. As mentioned in the Methods, this allowed the CRyPTIC project to produce high-confidence MICs where two or more independent methods agreed on the value^6,11^, thereby reducing measurement error. We will therefore examine the effect of only using these high-confidence MICs.

The sensitivities of ten of the 11 antibiotics for which we have UKMYC data increased (clofazimine being the exception) compared to the Fails+Minors dataset, with increases ranging from +1.4% for rifampicin to +15.8% for linezolid (Fig. 1 and Table 2). Specificity was largely unchanged with only ethambutol (-2.2%) changing by more than an absolute percentage point. Only the increases for linezolid, amikacin and kanamycin were statistically significant; if we compare back to the Simple dataset then the increases for isoniazid, linezolid, moxifloxacin, levofloxacin, amikacin and kanamycin were statistically signficant. No changes in specificity were statistically significant. This suggests that measurement error is partly constraining the measured performance of the WHOv2 catalogue. Notably both rifampicin and isoniazid now achieve sensitivities and specificities above 95% which is a requirement to pass ISO 20776-2:2021^29^.

### Discrepants do not have lower growth but, for some drugs, the MICs lie near the ECOFF/ECV

Our starting hypothesis for why the genetics and pDST results remained different was that the discrepant samples were more likely to have poor bacterial growth on the UKMYC plates after two weeks incubation as this could lead to measurement error. Examining the distributions of bacterial growth (averaged from the two positive control wells) showed that this is not true (Fig. S2 & S3) as, whilst there was large variation between samples, there was no significant difference in the distributions between e.g. the True Positives (RR) and False Positives (RS). Whilst the original measurement was an MIC, this was binarised using an ECOFF/ECV^11^, and hence our next hypothesis was that discrepants could arise if their MICs were close to the ECOFF/ECV, again leading to measurement error. Results varied by drug (Fig. S2 & S3), but this appears at least partly true for ethambutol, moxifloxacin and delamanid. It is clear that the less bimodal the MIC distribution, the more likely this effect will lead to misclassification and thereby discrepants.

### The performance of gnomonicus is similar to TB-Profiler

The Diverse Testset of 2,663 samples was also run through `TB-Profiler` (Methods). The predicted effects of an antibiotic on a sample were compared where (i) both methods returned a result for a sample and (ii) there was also a matching pDST result as per Table 1. There were no significant differences between the sensitivities and specificities calculated using both `gnomonicus` and `TB-Profiler` (Fig. 2), however this approach is somewhat crude as it e.g. groups together `Susceptible` and `Unknown` predictions and does not consider differences in predictions at the level of an individual sample. Hence let us drill down into the differences between the two tools.

**Figure 2.**
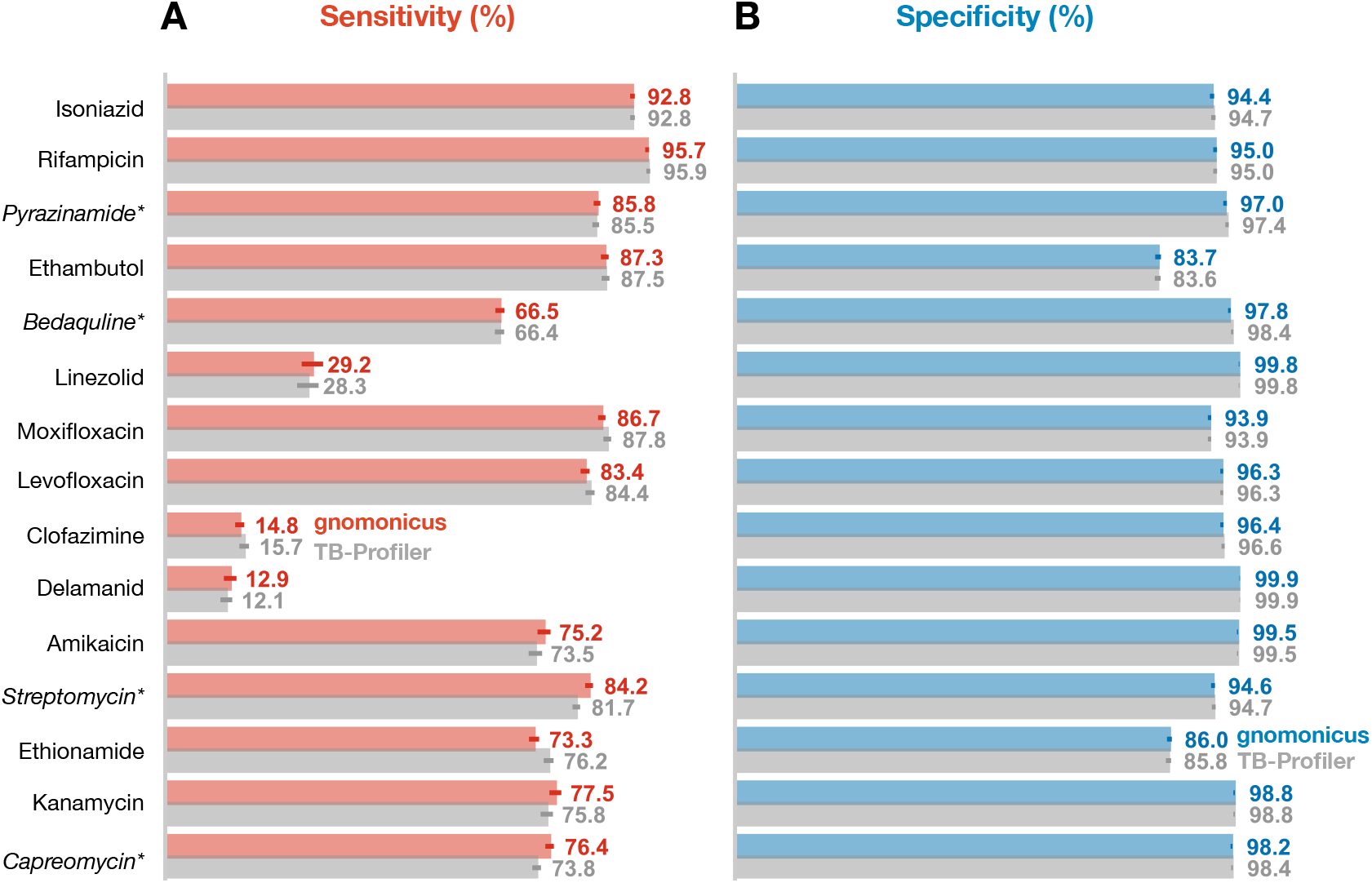
The performances for gnomonicus and `TB-Profiler` as measured using the 2,663 samples of the Diverse Testset are very similar. The (**A**) sensitivities and (**B**) specificities for all the drugs in WHOv2 are reported. To enable comparison for all drugs, the gnomonicus results are the Fail+Minors implementation. Four drugs (marked with an asterisk) have phenotypes from MGIT testing. Applying a McNemar test to the paired results suggests none of the difference reach significance (*p <* 0.05).

Two methodological differences become apparent when comparing the predictions in aggregate across all drugs (Table 3): firstly only `gnomonicus` returns a result of `Fail` (n=27) and secondly `gnomonicus` returns nine times more `Unknown` results (n=1,071 v 115). The former classification is unique to `gnomonicus` whilst the latter is due to our decision to return a result of `Unknown` for mutations in resistance genes that are not explicitly listed in WHOv2. In these cases `TB-Profiler` would instead return a result of `Susceptible`. The aim of both classifications is to avoid Very Major Errors (VMEs); these are defined as resistant samples incorrectly predicted as susceptible^30^. The tools `gnomonicus` and `TB-Profiler` make 881 and 1,096 VMEs, respectively. The `Fail` category avoids five VMEs on this dataset whilst the additional `Unknown` calls made by `gnomonicus` accounts for 205. Both tools (and thence the WHOv2 catalogue) have an aggregate VME rate far in excess of the 3% required by the CLSI (or 1.5% mandated by the FDA) but this is mainly driven by the poor performance of a few drugs (Fig. 2, Table S3).

**Table 3:** The majority of classifications made by `gnomonicus` and `TB-Profiler` agree (12,729/13,821, 92.1%). Most of the differences (965/13,821, 7.0%) occur when `gnomonicus` returns an `Unknown` result due to a mutation in a resistance gene that is not listed in WHOv2 whilst `TB-Profiler` returns `Susceptible`. A minority of classifications (63/13,821, 0.46%) are true discrepants in the sense that one method returns `Resistant` and the other `Susceptible`. The number of classifications that are measured as resistant by a pDST method are given in parentheses. Table S3 contains a detailed breakdown by drug.

|  |  | TB-Profiler |  |  |  |
| --- | --- | --- | --- | --- | --- |
|  |  | Resistant | Susceptible | Unknown | Total |
| gnomonicus | Resistant | 3,705<br>(3,297) | 43<br>(33) | 0<br>(0) | 3,748<br>(3,330) |
|  | Susceptible | 20<br>(18) | 8,932<br>(853) | 23<br>(10) | 8,975<br>(881) |
|  | Unknown | 14<br>(11) | 965<br>(205) | 92<br>(33) | 1,071<br>(249) |
|  | Fail | 4<br>(3) | 23<br>(5) | 0<br>(0) | 27<br>(8) |
|  | Total | 3,743<br>(3,329) | 9,963<br>(1,096) | 115<br>(43) | 13,821<br>(4,468) |

Overall both tools agree on a classification of `Resistant`, `Susceptible` or `Unknown` in 92.1% (12,729/13,821) of cases. As noted above, the majority of the difference (7.0%, 965/13,821) is due to our decision to return a result of `Unknown` for mutations in resistance genes not listed in WHOv2. That leaves 127 discrepants (0.92 %); about half (64) of which should not lead to clinical error as one or other of the tools is returning a result of `Unknown` or `Fail` which should, in practice, trigger additional investigations or tests. That leaves 63 (0.46%) predictions as true discrepants where one method has returned a result of `Resistant` and the other `Susceptible`.

The discrepants split into two groups; in 43 samples `gnomonicus` predicts resistance whilst `TB-Profiler` does not and in the remaining 20 samples the situation is reversed with the latter group being more enriched for resistance (18/20, 90.0% v 33/43, 76.7%). Of these 63 predictions, `gnomonicus` and `TB-Profiler` are therefore correct in 33+2=35 cases and 18+10=28 cases, respectively, and hence the behaviour of both tools is not significantly different on this dataset (McNemar test, p-value 0.45). Not all of these predictions are independent as there are several members of the fluoroquinolones and aminoglycosides in WHOv2 which all have similar resistance mechanisms. Only considering a single member from each drug class does not alter the result.

Breaking down the 63 discrepants by drug shows that all drugs have between one (rifampicin, linezolid and delamanid) and nine (capreomycin) discrepant predictions (Table 4). All the genetic variants driving these 63 discrepant predictions are given in the Supplement (Table S4). The main conclusion is that the discrepants are mainly due to differences the upstream genetic processing pipelines detecting different genetic variants in a small number of samples rather than how the WHOv2 catalogue has been translated and incorporated into both tools. This is encouraging but illustrates the importance of the upstream variant caller. The only cases where a variant is detected by both tools but only one predicts resistance is where there is evidence of two SNPs at a single locus in a resistance gene, here *gyrA* (three samples, Table S4) and *katG* (one sample). `TB-Profiler` identifies the presence of two mutations (e.g. D94N and D94Y in *gyrA*) and since both are associated with resistance to levofloxacin and moxifloxacin predicts resistance to both drugs. Since all seven cases are phenotypically resistant, `TB-Profiler` is correct. By contrast, `gnomonicus` notes that there is a heterogeneous call but does not resolve it further and therefore no catalogue rules are triggered and an incorrect result of susceptible is returned. This accounts for seven of the 20 cases where `TB-Profiler` calls resistance and `gnomonicus` does not (Table S4). Five of the remaining 13 cases are due to `TB-Profiler` detecting a single base deletion in *Rv0678 / mmpR5* at position 198.

**Table 4:** The 63 discrepant classifications produced by `gnomonicus` and `TB-Profiler` broken down by drug. The quadrants from the truth table are listed with the `gnomonicus` result first and the `TB-Profiler` result second. The numbers of paired discrepants are too small to support useful statistical testing.

| Drug | RR | RS | SR | SS |
| --- | --- | --- | --- | --- |
| Isoniazid | 483 | 2 | 1 | 467 |
| Rifampicin | 480 | 0 | 1 | 497 |
| Pyrazinamide | 256 | 3 | 0 | 351 |
| Ethambutol | 360 | 0 | 2 | 576 |
| Bedaquiline | 268 | 5 | 3 | 456 |
| Linezolid | 34 | 1 | 0 | 882 |
| Moxifloxacin | 290 | 1 | 4 | 632 |
| Levofloxacin | 290 | 1 | 4 | 636 |
| Clofazimine | 64 | 2 | 3 | 878 |
| Delamanid | 18 | 1 | 0 | 739 |
| Amikacin | 213 | 5 | 0 | 719 |
| Streptomycin | 224 | 6 | 0 | 313 |
| Ethionamide | 304 | 4 | 0 | 498 |
| Kanamycin | 226 | 5 | 0 | 704 |
| Capreomycin | 195 | 7 | 2 | 584 |
| Total | 3,705 | 43 | 20 | 8,932 |

Twenty two of the 43 cases where `gnomonicus` predicts resistance are due to variants identified in the *rrs* gene. Only 17 samples are affected because mutations in this gene are implicated in aminoglycoside resistance but of these nine are due to the presence of a1401g at varying levels of support which is not detected and therefore called by `TB-Profiler`. Since 15 out of 16 of the resulting predictions are phenotypically resistant, we infer that `gnomonicus` is correct in these cases. Another five cases are due to other variants detected in *rrs* by `gnomonicus` and interestingly, four cases are due to the identification of large deletions in *ddn* (22 bases), *Rv0678* (76 bases) *gid* (133 bases) and *ethA* (700 bases) leading to frameshifts and/or loss of function not picked up by `TB-Profiler`.

## Discussion

We have characterised the performance of our antimicrobial resistance prediction tool, `gnomonicus`, using the second edition of the WHO catalogue of resistance-associated mutations^8^ and compared its performance to that of `TB-Profiler`^9^, an established bioinformatic tool with similar functionality. Whilst WHOv2 is an accepted and well-known catalogue we have had to, in effect, translate the catalogue since the published catalogue is not computer-parsable and therefore we are testing two distinct statements: (i) that our translation is correct and (ii) that the reported performance for WHOv2^8^ is representative of what would be expected in the clinic. Unfortunately the samples that make up the WHOv2 Training Dataset are not (yet) publicly available which would have helped us with the first point. There are likely very good reasons for this, for example data owners may be willing to share samples with the WHO but not the public, but it does prevent researchers, such as ourselves, from reproducing the analysis that led to WHOv2 which is important to find errors and gain trust. We cannot therefore formally disentangle whether the observed differences are due to differences between the WHO Training Datatset and our publicly-available Diverse Testset, errors we have made parsing the catalogue, or assembly and variant calling differences. Some reassurance is gained on both points from `gnomonicus` (i) achieving similar performance for many of the 15 antibiotics covered by WHOv2 using our Diverse Testset of 2,663 *M. tuberculosis* samples and (ii) that its results are in close agreement with those produced by `TB-Profiler`^9^. If we only compare our results to high-confidence pDST results, the two main antitubercular drugs, rifampicin and isoniazid, achieve sensitivities and specificities above 95% which is required by ISO 20776-2:2021^29^.

Since there is no published guidance on how to *apply* WHOv2 to a WGS sample we have chosen to make three enhancements; the first is to explicitly flag genetic loci which are associated with resistance when there are insufficient reads to identify the allele. These we call `Fails` and, whilst small in number (2.5% of samples had at least one `Fail`), they correlate with poor sequencing quality and usually indicate a sample needs to be re-sequenced. The second improvement is we have chosen to classify mutations in genes known to be associated with resistance as `Unknown` even if they are not explicitly listed in the WHOv2 catalogue. This is possible because our GARC grammar (defined in the Supplement) allows wildcards such that our catalogues can contain a single rule encoding logic like “any missense mutation in the coding region of gene X” which, in addition to a way of prioritising rules, is necessary to enable this functionality. Finally, `gnomonicus` identifies any resistance-associated variant if it is supported by at least three (short) reads, thereby allowing minor alleles to contribute to resistance prediction. This has been shown to boost the sensitivity for fluoroquinolones^31^, rifampicin^32^ and bedaquline^33^ and the WHOv2 report also showed how lowering the fraction of reads supporting a variant call often increased the sensitivity^8^. The net effect of these changes was that nine antibiotics saw an increase in sensitivity of more than two absolute percentage points but this was only statistically significant for bedaquline, clofazimine and capreomycin; a larger dataset is needed to draw definite conclusions. Only two drugs saw a drop in specificity of more than one absolute percentage point and neither decrease was statistically significant. The effect of these enhancements can also be clearly seen when the results of `gnomonicus` and `TB-Profiler` are compared; the latter only implements the last enhancement and therefore does not return a result of `Fail` and `gnomonicus` returns almost an order of magnitude more `Unknown` results due to the second enhancement, both of which help reduce the number of Very Major Errors.

To calculate the sensitivity and specificity one must have a binary outcome yet the WHOv2 catalogue yields three: `Resistant`, `Susceptible` and `Unknown`. This was achieved in the WHOv2 report by assuming all samples classified as `Unknown` were susceptible. So that we may compare our results to those reported for WHOv2^8^ we have done the same (Fig. 1, 2 & Table 2). As we have noted `gnomonicus` returns more `Unknown` classifications than `TB-Profiler`, mainly because it detects and calls any SNP, insertion or deletion in a resistance gene that is not explicitly listed in WHOv2: combining `Susceptible` and `Unknown` results therefore masks this difference. Aggregating `Susceptible` and `Unknown` results like this is reasonable for antibiotics where the majority of the genetic determinants have been discovered since the conditional probability of a novel variant conferring resistance is low, but for drugs like bedaquiline or even pyrazinamide this assumption breaks down. For these drugs at least, it seems sensible to report an `Unknown` result for the drug, rather than assume it is `Susceptible`. Other studies have shown how, in some cases, one can take advantage of the correlations between different drugs to infer that they are likely `Susceptible`^34^.

Most catalogues of resistance-associated variants and bioinformatic tools, such as `TB-Profiler` use a three-valued logic: `Resistant`, `Susceptible` and `Unknown`. Here we introduce a fourth value, `Fail`, that is different to the others since it indicates we are lacking information at one or more genetic loci associated with resistance; in practice this usually means the sequencing was of poor quality and the run needs to be repeated. It is worth reflecting, however, on the `Unknown` value as this is, in fact, composed of two distinct cases. These are delineated by whether the genetic variant has been seen in sufficient clinical samples to have adequate statistical support or not. If not then the label is transitory; collecting more samples will improve the statistics and its effect will become associated with a definite label – we suggest this retains the `Unknown` label. But if sufficient samples are present in the Training Dataset then it corresponds to genetic variants, such as M306V/I in *embB*, where the minimum inhibitory concentration distribution straddles the ECOFF/ECV. Collecting more samples changes nothing and one can argue another *definite* value, distinct from `Resistant` or `Susceptible`, is therefore needed. Naming such a value is controversial (e.g. `Intermediate`) but its existence is unavoidable and undeniable.

Clearly it would have been preferable to be able to isolate our implementation of the WHOv2 catalogue in our Mycobacterial pipeline from the performance of the WHOv2 catalogue but this was not possible due to the lack of publicly available datasets. Also, whilst we have described our 2,663 samples as a Diverse Testset it is likely not truly independent since at least the 1,000 UKMYC samples form part of the WHO Training Dataset. Other shortcomings of our Diverse Testset include that (i) we were unable to obtain for all drugs a 50:50 split in resistance/susceptibility which reduces our statistical resolving power for those drugs and (ii) that it is not designed to be genetically diverse so does not test all rules in a catalogue. Finally, despite our best efforts, there are insufficient samples resistant to bedaquline, linezolid and delamanid in the Diverse Testset; additional, targeted collection of samples which are then made publicly available is the only answer here.

Our aspiration is that open-source tools, like `gnomonicus`, will help accelerate the uptake of whole-genome sequencing by public health bodies for tuberculosis surveillance and diagnostics^35^. To characterise its performance we have constructed and made available a Diverse Testset of 2,663 *M. tuberculosis* samples; we hope this will be of use to other researchers and could become a standard dataset upon which new tools and/or catalogues could be tested, allowing the consistent reporting of preliminary performance. All samples and their results can be downloaded using the attendant GitHub repository^1^ which also contains code allowing all analysis to be repeated and all figures to be redrawn.

## Acknowledgements

We are grateful to Jody Phelan for helpful discussions and to EIT Oxford for deploying our pipeline in their cloud platform and to ORACLE Corporation for access to their cloud.

## Ethics

All samples (both genetics and drug susceptibility data) were downloaded from public repositories. Ethics approval was previously obtained by the CRyPTIC project^6^.

## Author contributions

TEAP, DWC and PWF conceived of the design. JW, CB, MB, MC, BC, AH, JK, MLA, RS, HT, TEAP, DWC, RT and PWF built and tested the Mycobacterial pipeline. SS and KG contributed to the co-ordination and development of EIT GPAS software.

