## Supplemental Information for "Characterising the performance of an antibiotic resistance prediction tool, gnomonicus, using a diverse testset of 2,663 *Mycobacterium tuberculosis* samples"

### Contents

**1 Definition of the Grammar for Antimicrobial Resistance Catalogues (GARC) 3**

1.1 Catalogue Backus-Naur Form . . . . . . . . . . . . . . . . . . . . . . . . . . . . . 3

1.2 Prediction Backus-Naur Form . . . . . . . . . . . . . . . . . . . . . . . . . . . . . 5

**2 The `gnomonicus` software tool 6**

**3 The Mycobacteria Pipeline used by GPAS 14**

S2 The discrepancies between the genetic prediction and the pDST result are not due to some samples growing less well on the UKMYC plates and therefore being more difficult to measure. 12

S3 The discrepancies between the genetic prediction and the pDST result are not due to some samples growing less well on the UKMYC plates and therefore being more difficult to measure (cont.). . . . . . . . . . . . . . . . . . . . . . . . . . . . . . . . . . . . . . . . . 13

S4 Mycobacterial Pipeline . . . . . . . . . . . . . . . . . . . . . . . . . . . . . . . . . . 15

### 1 Definition of the Grammar for Antimicrobial Resistance Catalogues (GARC)

The below are all described using standard Backus-Naur form (BNF)

#### 1.1 Catalogue Backus-Naur Form

This is a definition of the GARC grammar acceptable to use within a catalogue. Where <gene-name> is any valid gene or locus name (usually matching the regex [a-zA-Z0-9_]+) from the reference genome (usually a GenBank file).

Note that to ensure uniqueness it uses some non-standard terms. Since the wildcards mimic the BASH shell, * is reserved for 'any position' and hence the Stop codon is instead described via !. A missense mutation is ? and a synonymous mutation is described by a =. Since the underscore character can be found in gene names, it is inappropriate to use as a delimiter, and hence we instead use @. Hence the rule that specifies that if three or more reads are found in a sample that support a Ser315Thr mutation in the gene *katG* would be katG@S315T:3,R.

An important and fundamental difference to e.g. the WHOv2 catalogue is GARC allows for a hierarchy of rules; a single genetic variant can (and usually will) hit multiple rules with the most specific assumed to triumph. All genes have several generic rules that ensure that novel mutations in a resistance gene return a result of U, hence in the above example, the genetic variant will also have hit the rule katG@*?,U but since it also hit a more specific rule, that is the result that is returned. Rules can be logically combined with an AND which is represented as an ampersand (&); this is necessary to encode epistatic rules, as introduced in WHOv2. So they can be distinguished, amino acids are written in upper case whilst nucleotides are described in lower case. Two additional letters are reserved: Z (or z) for describing heterogenous calls and X (or x) for null calls i.e. where there are insufficient reads at a locus to make a definite call.

```
<complete-mutation> ::= <partial-mutation> | "^"<partial-mutation>
<partial-mutation> ::= <mutation> | <mutation>":"<number> | <mutation>":0."<number> | <partial-mutation>"&"<partial-mutation>
<mutation> ::=
               <gene-name>"@"<nucleotide><position><nucleotide> |
               <gene-name>"@"<amino-acid><number><amino-acid> |
               <gene-name>"@"<position>"\_ins\_"<nucleotides> |
               <gene-name>"@"<position>"\_ins\_"<number> |
               <gene-name>"@"<position>"_ins" |
               <gene-name>"@"<position>"\_del\_"<nucleotides> |
               <gene-name>"@"<position>"\_del\_"<number> |
               <gene-name>"@"<position>"_del" |
               <gene-name>"@"<position>"_indel" |
               <gene-name>"@"<position>"_fs" |
               <gene-name>"@"<pos-wildcard><wildcard> |
               <gene-name>"@"<nucleotide><pos>"?" |
               <gene-name>"@"<amino-acid><number>"?" |
               <gene-name>"@"<positive-position>"=" |
               <gene-name>"@del_0."<number> |
               <gene-name>"@del_1.0"

<wildcard> ::= "?" | "="

<positive-position> ::= <number> | "*"

<position> ::= <pos> | <pos-wildcard>

<pos-wildcard> ::= "*" | "-*"

<pos> ::= <number> | "-"<number>

<nucleotides> ::= <nucleotide> | <nucleotide><nucleotide>

<nucleotide> ::= "a" | "c" | "t" | "g" | "x" | "z"

<amino-acid> ::= "A" | "C" | "D" | "E" | "F" | "G" | "H" | "I" | "K" | "L" | "M" | "N" | "O" | "P" | "Q" | "R" | "S" | "T" | "V" | "W" | "X" | "Y" | "Z" | "!"

<number> ::= "0" | "1" | "2" | "3" | "4" | "5" | "6" | "7" | "8" | "9" | <number><number>
```

#### 1.2 Prediction Backus-Naur Form

Due to wildcards not being intended for use for prediction, the grammar for prediction is slightly simplified to reflect this. As before, <gene-name> is still any valid gene or locus name (usually matching the regex [a-zA-Z0-9_]+)

```
<complete-mutation> ::= <mutation> | <mutation>":"<number> | <mutation>":0."<number> | <complete-mutation>"&"<complete-mutation>
<mutation> ::=
               <gene-name>"@"<nucleotide><pos><nucleotide> |
               <gene-name>"@"<amino-acid><number><amino-acid> |
               <gene-name>"@"<pos>"\_ins\_"<nucleotides> |
               <gene-name>"@"<pos>"\_ins\_"<number> |
               <gene-name>"@"<pos>"_ins" |
               <gene-name>"@"<pos>"\_del\_"<nucleotides> |
               <gene-name>"@"<pos>"\_del\_"<number> |
               <gene-name>"@"<pos>"_del" |
               <gene-name>"@"<pos>"_indel" |
               <gene-name>"@"<pos>"_fs" |
               <gene-name>"@del_0."<number> |
               <gene-name>"@del_1.0"

<pos> ::= <number> | "-"<number>

<nucleotides> ::= <nucleotide> | <nucleotide><nucleotide>

<nucleotide> ::= "a" | "c" | "t" | "g" | "x" | "z"

<amino-acid> ::= "A" | "C" | "D" | "E" | "F" | "G" | "H" | "I" | "K" | "L" | "M" | "N" | "O" | "P" | "Q" | "R" | "S" | "T" | "V" | "W" | "X" | "Y" | "Z" | "!"

<number> ::= "0" | "1" | "2" | "3" | "4" | "5" | "6" | "7" | "8" | "9" | <number><number>
```

#### 2 The gnomonicus software tool

The second edition of the WHO catalogue of resistance-associated mutations (WHOv2)[1] was translated into our Catalogue GARC grammar (formally defined in §1.1) using some bespoke Python code[2]. The resulting single CSV file is publicly available in a GitHub repository[3] that also contains other catalogues, including WHOv1. Our implementation of WHOv2 deviates slightly from the original catalogue; the majority of these were required to translate the catalogue into GARC. A key point of difference is our inclusion of a 'Failed' prediction. These will be triggered if there are insufficient genetic reads (a 'null call') at a loci known to be associated with resistance; this can be either a SNP or a insertion/deletion, like a frameshift. The variant caller used by the pipeline deployed in GPAS defines a null call as any loci having only two, one or zero reads at a locus. The other main difference is we include so-called 'wild-card' rules for resistance genes such that any mutation in that gene will return a result of 'Unknown' if it is not classified by the catalogue as either 'Resistant' or 'Susceptible'.

`gnomonicus` is a Python package[4] and can be installed either via the Python Packaging Index (PyPi) * or directly via its GitHub repository †. It is freely-available for research use but is not licensed for commercial use. It ingests

1. a catalogue in the GARC format as described above[3]
2. a GenBank file for the reference genome that used when constructing the variant call file (VCF); here the reference is version 3 of the H37Rv reference genome (`NC_000962.3`).
3. a VCF file describing the variation (or otherwise) in the sample with respect to the above reference.

One can specify via a command-line flag if `gnomonicus` should only consider and report on the resistance genes, as defined by the supplied catalogue, or whether mutations should be reported for all genes. It returns either a single JSON object describing the detected mutations and their predicted effects according to the supplied catalogue and/or a series of CSV files. The former is more useful when using on a single sample whilst the latter is particularly useful when using `gnomonicus` to report on all detected mutations in all genes as it minimises the disc space required to store the outputs and means they can be trivially aggregated. Note that, as is common in tuberculosis genetics, we assume the 100 non-coding bases upstream of the start of a gene form the promoter. `gnomonicus` has several dependencies, including `piezo` ‡ and `grumpy` § – the latter being a reimplementation of `gumpy` ¶ in Rust to improve performance.

---

*`https://pypi.org/project/gnomonicus/`
†`https://github.com/oxfordmmm/gnomonicus`
‡`https://github.com/oxfordmmm/piezo`
§`https://github.com/oxfordmmm/grumpy`
¶`https://github.com/oxfordmmm/gumpy`

| Number of drug phenotypes | Number of samples |
|---|---|
| 1 | 347 |
| 2 | 702 |
| 3 | 581 |
| 4 | 33 |
| Total | 1,663 |

Table S1: The distribution of number of phenotypes in the 1,663 samples with MGIT phenotypes

| | Simple | Nulls+Minors | Nulls+Minor (High confidence) |
|---|---|---|---|
| Isoniazid | 95.2 ±0.4 | 94.4 ±0.4 | 95.3 ±0.3 |
| Rifampicin | 95.6 ±0.4 | 94.7 ±0.4 | 95.5 ±0.4 |
| Pyrazinamide* | 96.4 ±0.5 | 96.1 ±0.5 | |
| Ethambutol | 69.3 ±1.0 | 68.5 ±1.0 | 67.4 ±0.9 |
| Bedaquiline* | 96.8 ±0.5 | 97.0 ±0.3 | |
| Linezolid | 95.9 ±1.8 | 94.2 ±1.6 | 96.1 ±1.2 |
| Moxifloxacin | 86.0 ±0.9 | 85.3 ±0.9 | 86.2 ±0.8 |
| Levofloxacin | 92.0 ±0.7 | 91.6 ±0.7 | 92.5 ±0.6 |
| Clofazimine | 57.9 ±3.4 | 61.8 ±2.4 | 31.6 ±2.8 |
| Delamanid | 93.7 ±3.0 | 94.0 ±2.9 | 100.0 ±0.0 |
| Amikacin | 98.6 ±0.3 | 98.2 ±0.4 | 98.1 ±0.4 |
| Streptomycin* | 92.8 ±0.6 | 91.6 ±0.6 | |
| Ethionamide | 69.8 ±0.9 | 68.3 ±0.9 | 68.9 ±0.9 |
| Kanamycin | 96.8 ±0.4 | 96.2 ±0.5 | 95.8 ±0.4 |
| Capreomycin* | 94.3 ±0.6 | 95.1 ±0.5 | |

Table S2: (Related to Table 2) How the Positive Prediction Value (%) changes as we first allow null calls and minor alleles to contribute and then only consider samples with high-confidence MICs. The latter is only possible for those drugs with phenotypes derived from the UKMYC plates and hence there are no values for pyrazinamide, bedaquline, streptomycin and capreomycin.

| Drug | pDST | gnomonicus \| TB-Profiler prediction RR | RS | FR | FS | SR | SS | SU | UR | US | UU |
|---|---|---|---|---|---|---|---|---|---|---|---|
| Isoniazid | R | 458 | 1 | 3 | 0 | 1 | 24 | 1 | 0 | 8 | 3 |
| | S | 25 | 1 | 0 | 1 | 0 | 443 | 0 | 0 | 24 | 5 |
| Rifampicin | R | 453 | 0 | 0 | 0 | 1 | 20 | 0 | 0 | 1 | 0 |
| | S | 27 | 0 | 0 | 1 | 0 | 477 | 0 | 0 | 18 | 0 |
| Pyrazinamide | R | 247 | 2 | 0 | 0 | 0 | 28 | 0 | 1 | 7 | 6 |
| | S | 9 | 1 | 0 | 0 | 0 | 323 | 0 | 0 | 0 | 3 |
| Ethambutol | R | 249 | 0 | 0 | 1 | 1 | 22 | 3 | 0 | 4 | 7 |
| | S | 111 | 0 | 0 | 3 | 1 | 554 | 9 | 0 | 23 | 10 |
| Bedaquiline | R | 261 | 3 | 0 | 0 | 3 | 90 | 4 | 0 | 33 | 0 |
| | S | 7 | 2 | 0 | 0 | 0 | 366 | 0 | 0 | 5 | 0 |
| Linezolid | R | 32 | 1 | 0 | 0 | 0 | 76 | 0 | 0 | 8 | 0 |
| | S | 2 | 0 | 0 | 0 | 0 | 806 | 0 | 0 | 73 | 0 |
| Moxifloxacin | R | 247 | 1 | 0 | 0 | 4 | 29 | 1 | 0 | 5 | 0 |
| | S | 43 | 0 | 0 | 0 | 0 | 603 | 0 | 0 | 64 | 1 |
| Levofloxacin | R | 266 | 1 | 0 | 0 | 4 | 39 | 1 | 0 | 10 | 1 |
| | S | 24 | 0 | 0 | 1 | 0 | 597 | 0 | 0 | 54 | 0 |
| Clofazimine | R | 40 | 0 | 0 | 0 | 2 | 225 | 0 | 0 | 21 | 0 |
| | S | 24 | 2 | 0 | 0 | 1 | 653 | 0 | 0 | 30 | 0 |
| Delamanid | R | 17 | 1 | 0 | 0 | 0 | 85 | 0 | 0 | 37 | 0 |
| | S | 1 | 0 | 0 | 0 | 0 | 654 | 0 | 0 | 203 | 0 |
| Amikacin | R | 209 | 5 | 0 | 0 | 0 | 66 | 0 | 0 | 7 | 0 |
| | S | 4 | 0 | 0 | 1 | 0 | 653 | 0 | 0 | 53 | 0 |
| Streptomycin | R | 207 | 5 | 0 | 1 | 0 | 13 | 0 | 0 | 23 | 3 |
| | S | 17 | 1 | 1 | 6 | 0 | 300 | 0 | 0 | 36 | 1 |
| Ethionamide | R | 208 | 2 | 0 | 0 | 0 | 32 | 0 | 10 | 28 | 8 |
| | S | 96 | 2 | 0 | 1 | 0 | 466 | 4 | 3 | 104 | 34 |
| Kanamycin | R | 217 | 5 | 0 | 0 | 0 | 54 | 0 | 0 | 10 | 2 |
| | S | 9 | 0 | 0 | 1 | 0 | 650 | 0 | 0 | 45 | 5 |
| Capreomycin | R | 186 | 6 | 0 | 3 | 2 | 50 | 0 | 0 | 3 | 3 |
| | S | 9 | 1 | 0 | 3 | 0 | 534 | 0 | 0 | 28 | 0 |

Table S3: (Related to Table 3) Detailed breakdown for each drug in WHOv2 of the predictions made by `gnomonicus` and `TB-Profiler` and how they compare to the phenotypic drug susceptibility testing (pDST) result. The `gnomonicus` result is given first and the `TB-Profiler` second e.g. FS indicates that the former `Failed` that sample whilst the latter predicted it was `Susceptible`. Only combinations of results with finite numbers of samples are shown.

| | | | gnomonicus | | TB-Profiler | |
|---|---|---|---|---|---|---|
| | | | Gene | Mutation | Gene | Mutation |
| ENA run accession | Drug | pDST | | | | |
| ERR13286130 | BDQ | S | Rv0678 | 494_ins_ct:16 | | |
| ERR13289278 | BDQ | S | Rv0678 | 141_ins_c:11 | | |
| ERR2510311 | EMB | R | | | embB | p.Met306Leu |
| ERR2510328 | LEV | R | | | gyrA | p.Asp94Asn |
| | | R | | | gyrA | p.Asp94Tyr |
| | MXF | R | | | gyrA | p.Asp94Asn |
| | | R | | | gyrA | p.Asp94Tyr |
| ERR2510548 | INH | R | | | katG | p.Ser315Thr |
| | | R | | | katG | p.Ser315Asn |
| ERR2510654 | STM | R | rrs | c517t:4 | | |
| ERR2510725 | CAP | R | rrs | a1401g | | |
| ERR2510733 | CAP | S | rrs | a1401g | | |
| ERR2515255 | PZA | S | pncA | H71Y:5 | | |
| ERR2515622 | STM | S | gid | 161_del_133 | | |
| ERR2515758 | PZA | R | pncA | !187W | | |
| ERR3287358 | LEV | R | | | gyrA | p.Asp94Asn |
| | | R | | | gyrA | p.Asp94His |
| | MXF | R | | | gyrA | p.Asp94Asn |
| | | R | | | gyrA | p.Asp94His |
| ERR3287504 | LEV | R | | | gyrA | p.Asp94His |
| | MXF | R | | | gyrA | p.Asp94His |
| ERR4796390 | CAP | R | rrs | g1484t:5 | | |
| ERR4797003 | CAP | R | rrs | a1401g | | |
| ERR4797034 | AMI | R | rrs | a1401g:3 | | |
| | KAN | R | rrs | a1401g:3 | | |
| ERR4797106 | AMI | R | rrs | a1401g | | |
| | KAN | R | rrs | a1401g | | |
| ERR4797121 | CFZ | S | Rv0678 | V1L:9 | | |
| ERR4797458 | LZD | R | rrl | g2814t:11 | | |
| ERR4797640 | AMI | R | rrs | a1401g:4 | | |
| | KAN | R | rrs | a1401g:4 | | |
| ERR4799472 | CAP | R | rrs | a1401g:7 | | |
| ERR4799886 | CAP | R | rrs | a1401g:7 | | |
| ERR4810820 | LEV | R | gyrA | D94H:170 | | |
| | MXF | R | gyrA | D94H:170 | | |
| ERR4820776 | CAP | R | rrs | a1401g:5 | | |
| ERR4822595 | LEV | R | | | gyrA | p.Asp94Ala |
| | | R | | | gyrA | p.Asp94Gly |

| | | | | | | |
|---|---|---|---|---|---|---|
| | MXF | R | | | gyrA | p.Asp94Ala |
| | | R | | | gyrA | p.Asp94Gly |
| ERR4828223 | ETH | R | inhA | S94A | | |
| | INH | S | inhA | S94A | | |
| ERR4828998 | AMI | R | rrs | a1401g | | |
| | KAN | R | rrs | a1401g | | |
| ERR4829940 | DLM | R | ddn | 380_del_22 | | |
| ERR4829960 | ETH | S | ethA | 774_del_700 | | |
| ERR4829977 | EMB | S | | | embB | p.Gly406Asp |
| | ETH | R | fabG1 | c-15t:6 | | |
| | INH | R | fabG1 | c-15t:6 | | |
| ERR4830157 | AMI | R | rrs | a1401g:9 | | |
| | KAN | R | rrs | a1401g:9 | | |
| ERR4830291 | ETH | S | ethA | Q360!:16 | | |
| ERR4831167 | CFZ | S | Rv0678 | 140_ins_tc:5 | | |
| ERR4831746 | CFZ | R | | | mmpR5 | c.198delG |
| ERR4831769 | CFZ | S | | | mmpR5 | c.198delG |
| ERR5917669 | STM | R | rrs | a514c:3 | | |
| ERR5917741 | STM | R | rrs | a514c | | |
| ERR5917758 | STM | R | rrs | a514c | | |
| ERR8975663 | CFZ | R | | | mmpR5 | c.421_425delGATCTinsA |
| ERR8976047 | RIF | R | | | rpoB | p.His445Val |
| ERR9121747 | STM | R | rrs | c517t:4 | | |
| ERR9992658 | BDQ | R | Rv0678 | 383_del_1 | | |
| ERR9992730 | BDQ | R | Rv0678 | 66_del_76 | | |
| ERR9992895 | BDQ | R | | | mmpR5 | c.198delG |
| ERR9992913 | BDQ | R | | | mmpR5 | c.198delG |
| ERR9993051 | BDQ | R | | | mmpR5 | c.198delG |
| ERR9993193 | BDQ | R | Rv0678 | 138_ins_g:9 | | |
| SRR1144772 | PZA | R | pncA | L182S:13 | | |
| SRR1163087 | CAP | R | | | rrs | n.1401A>G |
| SRR1165525 | CAP | R | | | rrs | n.1401A>G |

Table S4: All the genetic variants responsible for the discrepant resistant predictions made by `gnomonicus` and `TB-Profiler`. The pDST result is shown to enable comparison. Note that both tools use slightly different nomenclature. As described in §1.1, the GARC grammar uses lower-case for nucleotides and upper case for amino acids, with ! reserved for Stop and ins/del to indicate an insertion or deletion. A colon indicates that the variant has less than 90% of reads (the default) supporting its call; in these cases the number of reads in support is given after the colon with three being the minimum to avoid false positive calls. `TB-Profiler` uses the HGVS nomenclature.

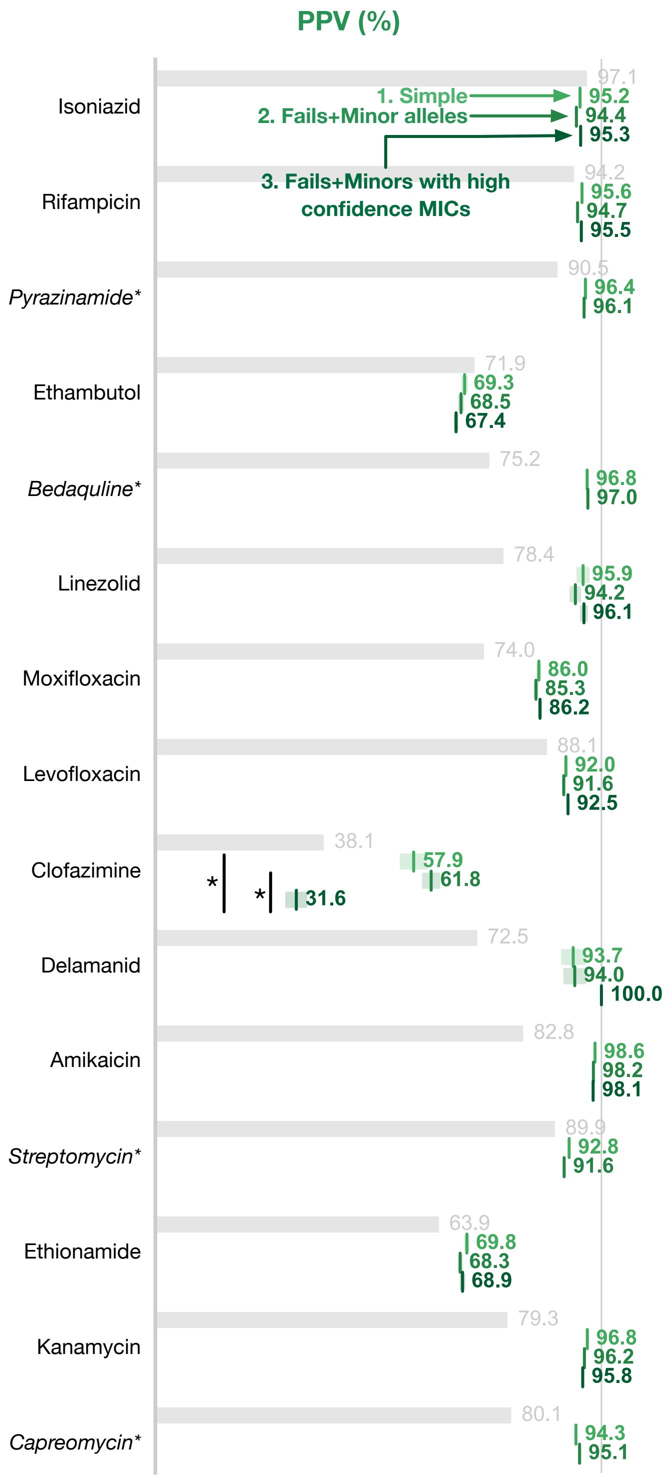

Figure S1: (Related to Figure 1) The `gnomonicus` AMR tool produces comparable AMR prediction performance, as measured by the positive predictive values, on the 2,663 validation samples as for reported for the WHOv2 catalogue. The grey bars are the performance reported on the WHO Training Set and since are calculated on a different, larger dataset are not directly comparable. Four drugs (marked with an asterisk) have phenotypes from MGIT testing. Three values are reported for the Mycobacterial Genetics Pipeline: (1) a simple translational of the WHOv2 catalogue, (2) then allowing resistant minor alleles with three or more reads to be called and, lastly, (3) only comparing to high confidence MICs in addition to calling minor alleles. Since the latter can only be calculated for the antibiotics with UKMYC MICs, no values can be reported for the four drugs measured by MGIT

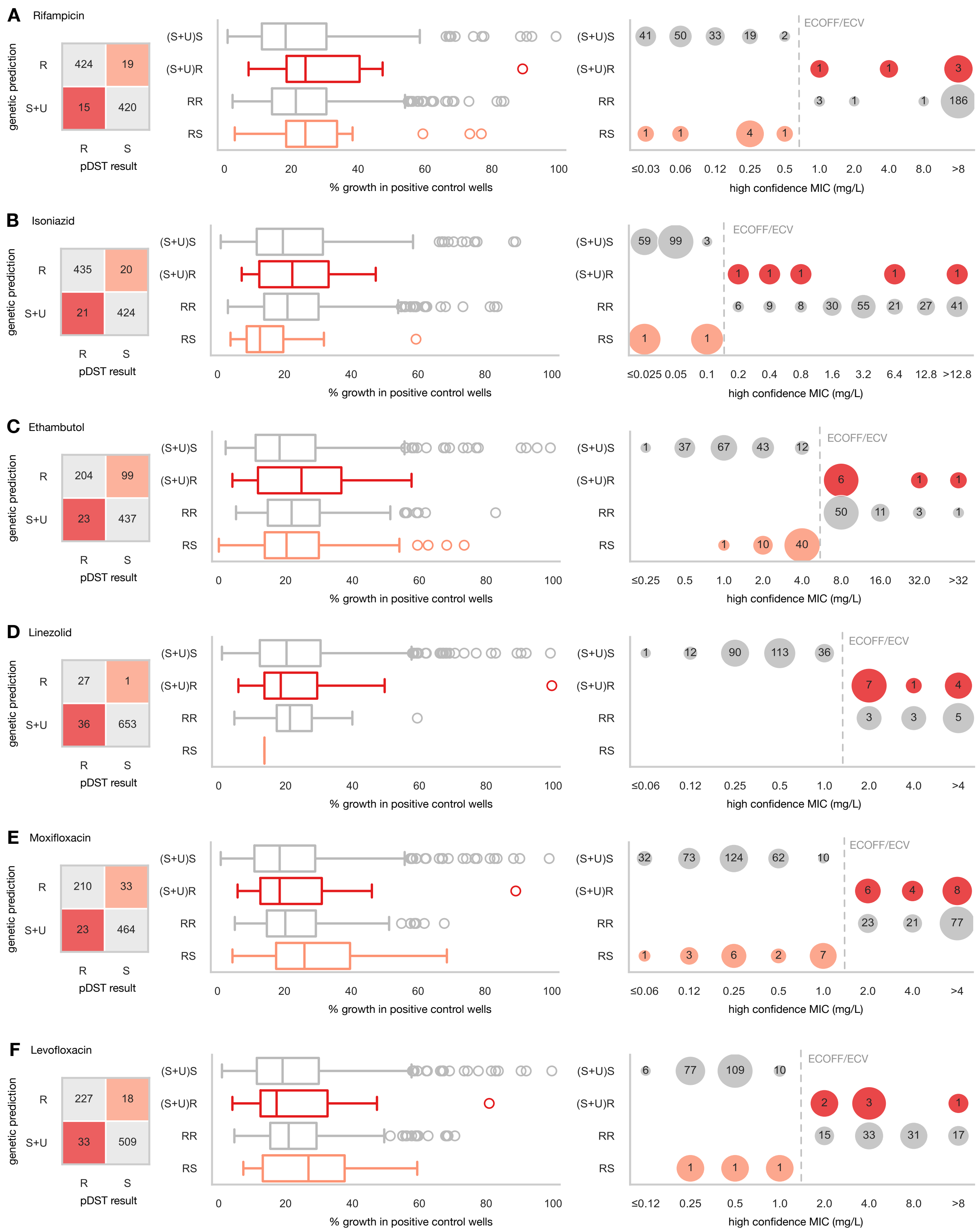

Figure S2: The discrepancies between the genetic prediction and the pDST result are not due to some samples growing less well on the UKMYC plates and therefore being more difficult to measure. For some drugs, one or both of the discrepant classes, are more likely to have an MIC close to the ECOFF/ECV whilst for some other drugs it appears not all the genetic basis for resistance is yet understood. All genetic predictions used the Fails+Minor Alleles implementation of the WHOv2 catalogue and only high confidence MICs were used. MIC distributions are shown for just one of the two 96-well plate designs used. Abbreviations: (S+U)S predicted sensitive or unclassified, pDST sensitive; (S+U)R predicted sensitive or unclassified, pDST resistant, RR predicted resistant, pDST resistant, RS predicted resistant, pDST sensitive.

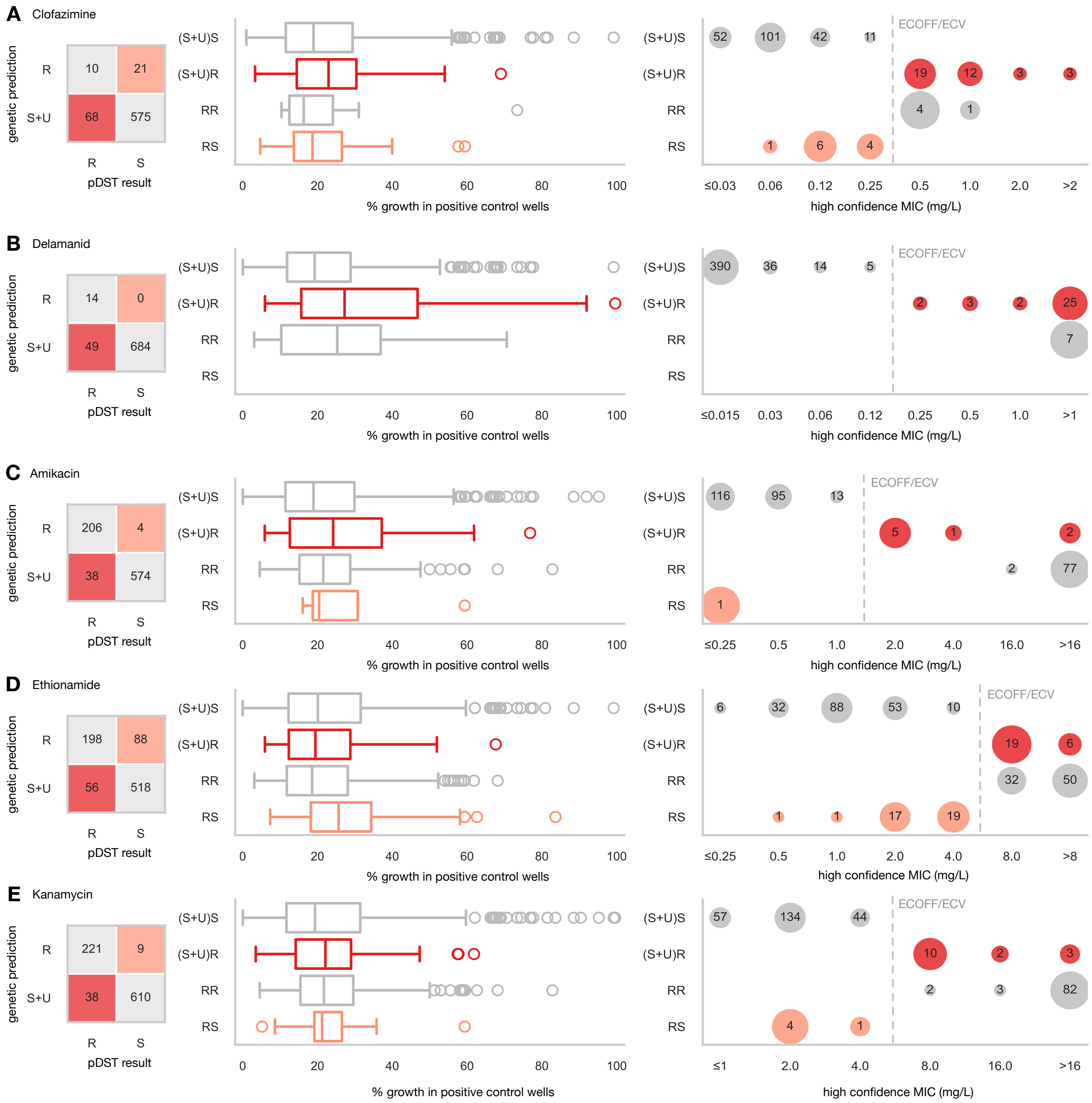

Figure S3: (Related to Figure S2) The discrepancies between the genetic prediction and the pDST result are not due to some samples growing less well on the UKMYC plates and therefore being more difficult to measure. For some drugs, one or both of the discrepant classes, are more likely to have an MIC close to the ECOFF/ECV whilst for some other drugs it appears not all the genetic basis for resistance is yet understood. All genetic predictions used the Fails+Minor Alleles implementation of the WHOv2 catalogue and only high confidence MICs were used. MIC distributions are shown for just one of the two 96-well plate designs used. Abbreviations: (S+U)S predicted sensitive or unclassified, pDST sensitive; (S+U)R predicted sensitive or unclassified, pDST resistant, RR predicted resistant, pDST resistant, RS predicted resistant, pDST sensitive.

#### 3 The Mycobacteria Pipeline used by GPAS

##### Overview

For completeness we describe in detail the Mycobacteria pipeline implemented in the EIT Global Pathogen Analysis Service that was used to process the FASTQ files of the 2,663 samples, producing the VCF files. The latter files are included in the attendant GitHub repository[5] along with instructions on how to apply our version of the WHOv2 catalogue using `gnomonicus` to these files to predict the effects of the 15 different drugs considered. The Mycobacteria pipeline (Figure S4) comprises a mixture of softwares written in a variety of languages. Nextflow was used throughout to standardise each of the pipeline steps and to manage the sequencing of tasks.

The pipeline begins with Human Read Removal (which . Subsequently, additional checks are made on the quality of the reads and only reads in the Mycobacteriacae family, and unknown reads, are retained. The species present is determined by competitively mapping the reads against a manifest of 186 manually-curated reference genomes of different Mycobacterial species. If the species belongs to a complex, then mykrobe is used to assign lineage (in the case of MBTC) or subspecies (e.g. in the case of *M. avium* complex). If the sample has sufficient reads belonging to the *Mycobacterium tuberculosis* complex then variants are called to inform antibiotic resistance prediction and the degree of relatedness between the sample and others in the database.

Data to be reported to the user is drawn together in the "Summary" step, which determines the full species name to report. Files generated by each of the steps of the pipeline are stored in buckets and can be downloaded by the user.

##### Remote Human Read Removal

Human read removal is the process of removing human reads from FASTQ files. As shown in Fig. S4, human read removal should also have already been carried out on the user's computer, with the remote human read removal step in the cloud providing added assurance. After this process is complete, the original uploaded files are deleted. Here we use `hostile`[6].

##### Read Quality Checks and Filtering

The next step checks the quality of the remaining reads and also applies some simple filtering as follows:

1. the FASTQ files are pre-processed with `fastp`[10]. This discards reads shorter than a specified threshold length: this is 50 bases for short-reads (Illumina) and 100 bases for long-reads (ONT). It also filters out for low-quality and low-complexity reads.

2. the reads are then classified by `kraken2` (v2.1.3) using the `Standard` index from June 2023[11]

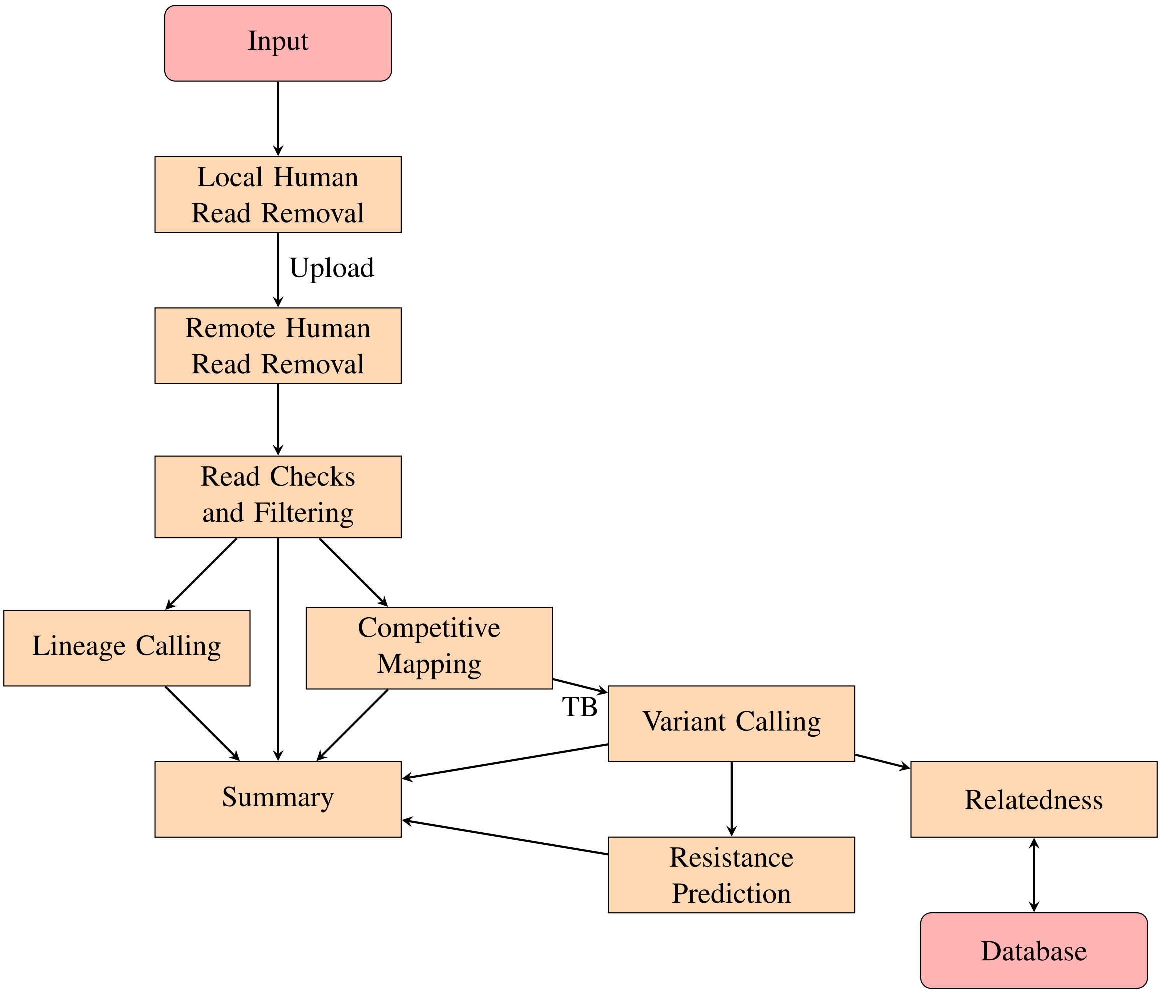

Figure S4: Mycobacterial Pipeline

3. The subset of reads classified as belonging to the Mycobacteriaceae taxon are retained, along unclassified reads.

Only samples with more than 10,000 reads (Illumina) or 1,000 reads (ONT) are progressed to subsequent steps.

#### Competitive Mapping

The aim of this step is to identify which Mycobacterial species are present in the filtered reads using a process we call 'competitive mapping'. This uses `minimap2` (v2.24-r1122)[7] to map reads against a curated collection of 186 reference mycobacterial genomes stored as a single multi-fasta file, called the manifest. `samtools` (v1.15.1)[9] is used to calculate coverage statistics against all reference genomes in the manifest. Competitive mapping outputs a number of statistics for each of the references; these are sorted by mean read depth (for Illumina this is essentially the genome with most reads, for ONT read length is accounted for). `minimap2` is run using the `-N 1000` command line flag to generate up to 1000 secondary alignments, and with `-x sr` and `-x map-ont`.

Where the number of reads mapped to *M. tuberculosis* exceeds a pre-determined threshold (which is different for Illumina and ONT), the sample is deemed to have sufficient reads to progress to genome determination. This happens even if *M. tuberculosis* is not the most prevalent species in the sample according to this step.

##### Lineage Calling

Competitive Mapping cannot resolve the differences between different lineages or different members of a species complex. Hence `mykrobe` (v0.13.0)[12] is used to predict the lineage and/or subspecies of any complexes present in the sample. The other capabilities of `mykrobe` are not used.

##### Variant Calling

`clockwork` (v0.12.3) is used for variant calling with short (usually Illumina) reads. `clockwork` is capable of several tasks, but here it is only used for variant calling and is configured not to remove unwanted reads or to trim reads. No mask is used. `clockwork` maps the reads against a provided reference, here version 3 of the H37Rv reference (`NC_000962.3`) using `minimap2`[7] to produce a pile-up. `samtools`[9] is used to remove PCR duplicates, outputing a sorted indexed BAM file. Variants are called independently by (i) `samtools`, which has a high sensitivity for SNPs, using `mpileup` command, and (ii) `cortex`[14], which has a lower sensitivity for SNPs but performs well for insertions and deletions. `minos`[15] adjudicates when there is a conflict. The minimum number of reads required for a call is three. The key output is a Variant Calling Format (VCF) file which lists the location of putative variations with respect to the reference genome and the evidence to support (or not) those variant calls. Rows containing null calls at loci associated with resistance are added using the VCF file generated by `samtools`.

##### Relatedness

Whilst determining if two samples are likely to epidemiologically related is 'simply' a matter of counting the number of SNP differences, comparing a single sample to a growing list of existing samples becomes ever more computationally challenging. We use an algorithm called `Find Neighbour 5`[‖] which calculates SNPs when given fixed length (i.e. no insertions) FASTA files. Each time a new sample is processed:

1. The FASTA file of the reference genome is read.

2. A mask is applied to ensure repetitive or highly variable regions are ignored.

3. The FASTA file of the sample is loaded and compared with the reference genome, base-by-base. If the sample is not identical to the reference at an unmasked base, the position is stored (along with a sample identifier).

4. These positions (arranged by nucleotide) are saved to disc.

[‖] `https://github.com/oxfordmmm/FN5`

5. The saves for each pre-existing sample are then loaded from disc, comparing SNP distance for each against the new sample. If the distance is above a user definable cutoff (20 for this pipeline), the distance calculation stops early (for efficiency).

6. The new sample is then added to the list of samples for future comparisons. This is performed in an asynchronous, distributed and thread-safe manner to ensure that distances are added appropriately.

##### Summary

The Summary task brings together data from other tasks in the pipeline and writes it into a single file. Some of the contents of this file are then copied into the database, which is used to populate the user interface and form the basis of reports downloaded by users. Data is typically unmodified by the Summary process, but for species identification the summary code draws together information from the Competitive Mapping and Lineage Calling steps to assign a species name and, as appropriate, a lineage or subspecies. The code also accommodates situations where mixed *M. tuberculosis* complex lineages are identified.
